# Destabilizing effects of eco-evolutionary feedback on community stability and the role of climate change

**DOI:** 10.1101/2025.06.04.657861

**Authors:** Qinghua Zhao, Frederik De Laender, György Barabas

## Abstract

What underpins the stability of natural systems is a central topic in ecology for decades. Evidence from simple communities with few species shows that eco-evolutionary feedback has the potential to stabilize community dynamics. Here, we show that these findings may not hold in more complex natural systems with many species. We first simulated an eco-evolutionary model of producer-herbivores dynamics, and found destabilizing, rather than stabilizing effect of the eco-evolutionary feedback (i.e. an increasing Lyapunov exponent). We then analyzed 11 long term monitoring data sets of natural aquatic communities, containing the temporal dynamics of a key individual-level trait (body mass) and of population densities. This analysis matched the model results and further showed that warming tends to make eco-evolutionary feedback even more destabilizing. Our results indicate that eco-evolutionary feedback may play a role in destabilizing community dynamics and the climate may mediate the magnitude.

## INTRODUCTION

In recent years, eco-evolutionary feedback, i.e. reciprocal interactions between populations dynamics and trait evolution, has been a main topic of study (Hendry 2017; De Meester *et al*. 2019; Govaert *et al*. 2019). While narrow sense eco-evolutionary feedback involves population-trait interactions within a single species, complex ecological relationships among species can make these feedback ripple through an entire community (Andrade-Domínguez *et al*. 2014; De Meester *et al*. 2019). Thus, eco-evolutionary feedback naturally extends to the community level (De Meester *et al*. 2019). For example, dominance of phytoplankton communities by toxic and nutritionally poor cyanobacteria can induce natural selection of tolerant *Daphnia* genotypes (Bernard *et al*. 1999; Sarnelle & Wilson 2005). This process in turn suppresses cyanobacteria growth (Schaffner *et al*. 2019) and lets non-cyanobacteria genotypes (e.g. green algae) flourish. While examples of such eco-evo feedback *sensu largo* are replete, it is unsure how they affect community stability. While there is theoretical evidence that eco-evolutionary feedback can be both stabilizing and destabilizing in small communities of simple architecture (Patel *et al*. 2018; Cortez *et al*. 2020), information is lacking for larger and more complex systems.

Anthropogenic climate change is expected to modify eco-evolutionary feedback (Barbour & Gibert 2021; Wood *et al*. 2021). Climate change manifests in multiple ways: temperature increase and deoxygenation are the two aspects (Schmidtko *et al*. 2017; Breitburg *et al*. 2018; Jane *et al*. 2021). Rising temperature and deoxygenation can, for example, change predator consumption rates and induce species selection on temperature (and deoxygenation) tolerance (Synodinos *et al*. 2021), thus altering extant ecological and/or evolutionary processes (Barbour & Gibert 2021; Jane *et al*. 2021). To date, there is lacking evidence that warming and deoxygenation would alter eco-evolutionary feedback, especially not in unconfined natural systems.

In this study, we first simulated producer-herbivore communities to numerically test the (de)stabilizing role of eco-evolutionary feedback. We quantified stability here as the dominant Lyapunov exponent (LE). Next, we attempted to compare these simulated results, by collating 11 aquatic datasets (ten food webs and one competitive community) spanning 10 to nearly 30 years of data on a key trait (body mass) and population densities. Then, to the 11 aquatic systems, we examined effect of temperature and deoxygenation on either stability or the magnitude of eco-evolutionary feedback. Overall, we found that eco-evolutionary feedback destabilizes community dynamics by increasing LE across both simulated and natural ecosystems. In natural systems, warming and deoxygenation made the eco-evolutionary feedback more destabilizing.

## METHODS

### Part 1. Model simulations

We simulated a community with multiple producer and herbivore species, by employing a general framework to describe the eco-evolutionary dynamics in the presence of intraspecific trait variation (Barabás & D’Andrea 2016) as,

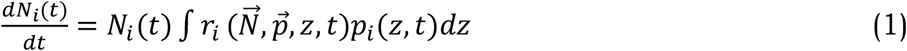

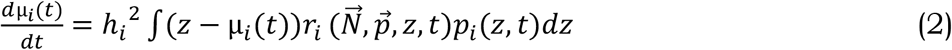

Where *N*_*i*_(*t*) is the population density of species *i* (both producers and herbivores) at time t, and μ_*i*_(*t*) is the mean trait value of species *i* at time t. The *p*_*i*_(*z, t*) is the distribution of species *i*’s trait value z at any moment of time *t*. The *h*_*i*_^2^ indicated heritability of species *i*, which is defined as the ratio of the additive and the total phenotypic variance.

Trait for individuals of a species is assumed to vary along a unidimensional axis. The phenotypic distribution of this trait is considered in the quantitative genetic limit. Under this limit, each of infinitely many loci contributes an infinitesimal additive value to the trait, on top of normally distributed environmental noise. In such assumptions, the distribution *p*_*i*_(*z, t*) of species *i*’s trait value z at any moment of time *t* is normally distributed, with phenotypic variance *σ*_*i*_^2^ that does not change in response to selection. Thus, one can recover the whole population density with its phenotype values, *N*_*i*_(*t*)*p*_*i*_(*z, t*), by just tracking the population density and mean trait value of each species in time.

To equation (1) and (2), once we know the detail of the per capita growth rate of species *i*’s phenotype z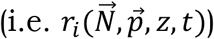, we can simulate the population abundance *N*_*i*_(*t*) and trait μ_*i*_(*t*) over time. Following (Barabás & D’Andrea 2016), the per capita growth rate can explicitly include two parts: its intrinsic growth rate and the effects of other individuals with phenotype *z*^’^. Thus the per capita growth rate 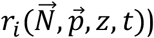 be written as

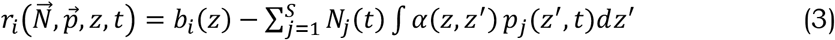

*b*_*i*_(*z*) is the intrinsic population growth of species *i*. S is number of interacting species. *α*(*z, z*^’^) is the interaction kernel, indicating the effect of the effect of another individual with phenotype *z*^’^ on the individual with phenotype z.

For producers, we consider *b*_*i*_(*z*) as a quadratic function (Barabás & D’Andrea 2016), *b*_*i*_(*z*) = 1 − *z*^2^/*θ*^2^, where *b*_*i*_(*z*) is positive on an interval [-θ, θ] but zero outside. Biological meaning of *b*_*i*_(*z*) is that any trait value falling outside the region [-θ, θ] is too extreme to efficiently forage for available resources and thus to get a positive growth rate. Then, the interaction kernel *α*(*z, z*^’^) for producers can also be dismantled into two parts: competition from another producer individual with phenotype *z*^’^ and consumption from herbivore individual with phenotype *z*^’^. The competitive interaction kernel is expressed as 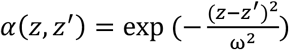 as (Barabás & D’Andrea 2016), with competition width ω. The more similar traits two individuals share, stronger competition. Following (Schreiber *et al*. 2011), the predatory interaction kernel is written as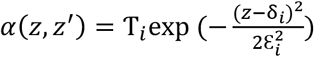, where herbivore’s attack rate T on prey *i* is maximal at an optimal trait value δ_*i*_ and decreases away from this optimal trait value. ε_*i*_ determines how steeply attack rate declines with distance from the optimal trait value δ_*i*_.

For herbivores, *b*_*i*_(*z*) is negative. We assume a larger trait value z has larger death rate, thus *b*_i_ (*z*) expressed as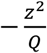. Here *Q* is the proportional constant. The predatory interaction kernel is same as the producers but times a conversion ecoefficiency *e*, written as *α*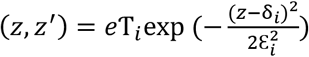, because only a fraction for energy is finally transfer to herbivores.

### Parameter setting

We generated a producer-herbivore community with 15 producers and 5 herbivores. Its connectance was set as 0.1, because the connectance is widely observed in natural systems (Dunne *et al*. 2002b, a). Next, heritability *h*_*i*_^2^ for the producer-herbivore system was generated from uniform distribution by function U[0.1, 0.4]. Competition width ω was drew from U[0.01, 0.05]. Intraspecific phenotypic standard deviations *σ*_*i*_ was generated from U[0.01, 0.1]. Initial trait for producers was produced by U[-1, 1], while herbivore’s initial trait was given as a larger values from U[0, 2]. Conversion ecoefficiency *e* was drew from U[0,1, 0.3]. Q set as a constant 100000. ε set as a constant 1, and optimal trait value δ drew from U[4, 8]. Attack rate T set as constant 0.1.

### Quantifying stability

For the producer-herbivore community (15 producers and 5 herbivores), we created 1000 replicates (parameter combinations) as described above. For each parameter setting, we calculated stability as dominant Lyapunov exponent *LE* as

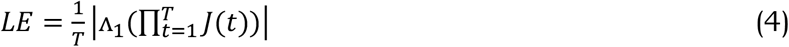

Here, *LE* is computed by taking the log absolute value of the dominant eigenvalue Λ_1_ of the sequential Jacobian product 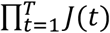, and then divided by time T. In perfect sense, the LE will converge in the limit as T approaches infinite. However, in practice, we only have finite time series, thus *LE* is calculated over the available length of Jacobian matrix *J*. The *J* can be written as

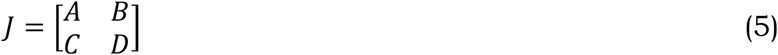

Where *A, B, C* or *D* is a block matrix. *A* and *B* are obtained by partial derivative of 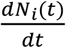 with respect to N(t) and μ(t), respectively. The block matrix *A* represents the effect of ecological process on itself (Eco→ Eco), and *B* represents the effect of trait evolution process on ecological process (Evo→ Eco). Similarly, the block matrix *C* and *D* are obtained by partial derivative of 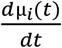 with respect to N(t) and μ(t), respectively. *C* represents the effect of trait evolution on ecological process (Evo→ Eco), and *D* represents the effect of trait evolution process on itself (Evo→ Evo). Considering the computation time, we didn’t use whole time series (0 to 100000) to compute the Jacobian matrix, but use a sampling time step of 10 (i.e. 0, 10, 20, ….., 1000000).

### Estimating the stability for ecological process or trait evolution

We first computed the stability (LE at eqn.4) for ecological process (i.e. abundance changes), by using the block matrix *A* (Eco→ Eco), because the *A* indicates the effect of ecological process on itself. Similarly, the stability (LE) for trait evolution was computed by only using the block matrix *D* (Evo→ Evo).

### Contribution of eco-evolutionary feedback on community stability

The eco-evolutionary feedback (Evo→ Eco→ Evo) includes two processes: (1) the effect of trait evolution on ecological process (Evo→ Eco), which is block matrix *B* in eqn 5; (2) the effect of ecological process on trait evolution (Eco→ Evo), which is block matrix *C* in eqn 5. If any elements of *B* and/or *C* within whole Jacobian *J* is nonzero, the eco-evolutionary feedback has contribution to whole community stability. Therefore, we arbitrarily set B and C in eqn 5 as zero (i.e. setting antidiagonal as zero), and then we computed the LE based on this revised *J*, called LE_0_. Finally, the magnitude of the eco-evolutionary feedback can be computed as the difference between original community stability (no elements set as zero in eqn 4) and LE_0_, i.e. LE-LE_0_. A positive value of it (LE-LE_0_) indicates the eco-evolutionary feedback destabilizes whole community stability, as it increased dominant Lyapunov exponent of whole community, and vice versa.

### Part 2. Empirical Data

The 11 aquatic ecosystems studied were Lake Monona (MO, 1995–2018), Lake Mendota (ME, 1995–2018), Lake Allequash (AL, 1982–2019), Lake Big Muskellunge (BM, 1990–2019), Lake Crystal Bog (CB, 1990–2016), Lake Crystal (CR, 1995–2019), Lake Sparkling (SP, 1990–2019), Lake Trout (TR, 1990–2019), Lake Oneida (ON, 1975–1995), Lake Zurich (ZU, 1982–2005), and Wadden sea (WS, 2001– 2010).

Each system reports three types of variables: (1) species abundance; (2) species trait (biomass or body-length); (3) water temperature. In addition, except for Lake Zurich and Wadden sea, the rest nine systems reported dissolved oxygen concentration. Phytoplankton of these system directly reported species as biomass, but zooplankton reported it as individual body-length. To be consistent trait across these systems, we transformed body-length into biomass. Specifically, individual body length L (mm) was transferred into biovolume (mm^3^) by 0.074 × *L*^2.92^ (Horn 1991), and its fresh body weight by multiplying the biovolume with a conversion factor of 1 (mg mm^-3^) (Havens 1995; Hwang & Heath 1999). The total fresh biomass (mg) of each zooplankton species was calculated by multiplying the individual biomass with its abundance. Note that fish species were excluded from analysis, because they were either not reported or yearly sampled only, and because the unit (catch per unit effort) of fish species changed over sampling time.

For consistency, each time series in each system was averaged to be triannual sampling frequency (if multiple sampling within a season, e.g. bimonthly or monthly sampling, they were averaged). Finally, each time series was scaled to zero mean and unit variance.

### Estimating species interactions and whole community stability

For each system above, we describe the changes in density of *k* species (*N*_1_, …, *N*_*k*_), and *l* population level evolving traits (e.g. biomass) with values *x*_1_, …, *x*_*l*_ as a general multispecies model. The general model is,

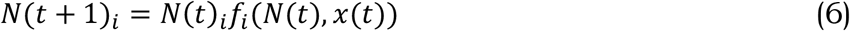

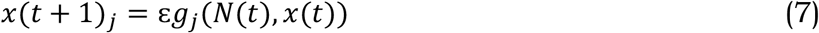

where *N*(*t*) = (*N*_1_(*t*), …, *N*_*k*_(*t*)) and *x*(*t*) = (*x*_1_(*t*), …, *x*_*l*_(*t*)). The function *f*_*i*_ stands for the per capita fitness of species *i*, which describes the growth of species *i* depending on the population densities of all species *N*(*t*) and all traits *x*(*t*). Similarly, the function *g*_*j*_ stands for the selection function for trait *j*, which describes how trait *j* evolves due to the selective pressures imposed by all species *N*(*t*) and all traits *x*(*t*). We make no initial assumptions about the exact form of *f*_*i*_ and *g*_*j*_. ε represents the timescale separation between the ecological and evolutionary dynamics. A small ε indicates that the evolutionary process occurs very slowly relative to the ecological processes (slow evolution), while a large ε indicates the evolutionary process occurs very quickly relative to the ecological processes (fast evolution). Notably, this model form is flexible to incorporate any combination of species interactions, e.g. herbivore-resource, mutualistic, and competitive interactions. In addition, this model form is also flexible to incorporate any number of evolving traits, e.g. single trait coevolution of multiple species and multi-trait evolution within one species.

With general equation (6) and (7) at hands, one can compute dominant Lyapunov exponent *LE* as eqn 4. Similarly, the Jacobian matrix *J* in eqn 4 can be written as eqn 5. Again, the block matrix *A* represents the effect of ecological process on itself (Eco→ Eco), and *B* represents the effect of trait evolution process on ecological process (Evo→ Eco). The block matrix *C* represents the effect of trait evolution on ecological process (Evo→ Eco), and *D* represents the effect of trait evolution process on itself (Evo→ Evo).

To estimate Jacobian matrix *J* for each of the 11 system, we employed the multiview distance regularised S-map (MDR S-map), because this approach had a higher accuracy to recover Jacobian than other techniques (Chang *et al*. 2021) and because and because the embedding dimension E in this study was smaller than the number of species (causal variables) in each of 11 systems. The MDR S-map is especially useful to reconstruct high-dimensional time-varying interaction networks for complex systems. It works by linking two methods (multiview embedding and regularised S-map), but shows a higher accuracy than each one alone. The formula and mechanism of MDR S-map see *Supplementary S1*.

Again, the stability (LE at eqn.5) for ecological process (i.e. abundance changes) was computed by using the block matrix *A* (Eco→ Eco), because the *A* indicates the effect of ecological process on itself. Similarly, the stability (LE) for trait evolution was computed by only using the block matrix *D* (Evo→ Evo). Finally, the magnitude of the eco-evolutionary feedback can be computed as the difference between original community stability (no elements set as zero in eqn 5) and LE_0_, i.e. LE-LE_0_, where LE_0_ was computed as by arbitrary setting B and C in eqn 5 as zero (i.e. setting antidiagonal as zero). Again, a positive value of it (LE-LE_0_) indicates the eco-evolutionary feedback destabilizes whole community stability, as it increased dominant Lyapunov exponent of whole community, and vice versa.

### Statistical analysis

To investigate if ecological process, trait evolution, eco-evolutionary feedback, or whole community dynamics is (de)stabilized over time, we should employ One-sample wilconxon test (*wilcox*.*test* function in R) to examine if their LEs is significantly different with 0. Since we conducted multiple comparisons, threshold for significance (*P*=0.05) might be lenient. Hence, we used *p*.*adjust* function to rectify all *P* values.

To investigate if ecological process, trait evolution, eco-evolutionary feedback, or whole community dynamics is (de)stabilized over time, we should employ One-sample wilconxon test (*wilcox*.*test* function in R) to examine if their LEs is significantly different with 0. Since we conducted multiple comparisons, threshold for significance (*P*=0.05) might be lenient. Hence, we used *p*.*adjust* function to rectify all *P* values.

Next, we employed linear regression to examine the relationship between warming and each of four stabilities above (i.e. LE for ecological process, LE for trait evolution, LE for eco-evolutionary feedback, and LE for whole community). *P* values were again rectified by *p*.*adjust* function. The warming rate (intensity of long-term warming) in each system was calculated by using the Theil–Sen median-based trend estimator, which is robust to episodic extreme events and provides a stable estimation for long-term trend even in the presence of outliers (Mohsin & Gough 2010; Chang *et al*. 2020). Similarly, oxygenation rate in each system was again estimated by using the Theil–Sen median-based trend estimator. A negative value of oxygenation rate indicates decline in aquatic oxygen content during studied period, and vice versa.

Deoxygenation rate of dissolved water oxygen concentration in each system was again calculated by using the Theil–Sen median-based trend estimator (Mohsin & Gough 2010; Chang *et al*. 2020).

Next, we examined the effect of warming rate on stability, because warming and deoxygenation showed no correlation in studied systems (Fig S3). We again used linear regression to examine the relationship between deoxygenation and each of four stabilities above. Again, *P* values were rectified by *p*.*adjust* function. Deoxygenation rate of dissolved water oxygen concentration in each system was again calculated by using the Theil–Sen median-based trend estimator (Mohsin & Gough 2010; Chang *et al*. 2020).

Finally, we employed linear regression to test the effect of warming rate on each of four stabilities above (i.e. LE for ecological process, LE for trait evolution, LE for eco-evolutionary feedback, and LE for whole community). We didn’t find warming had an association with oxygenation rate across 11 systems (Fig S3). *P* values were rectified by *p*.*adjust* function. Similarly, we used linear regression to examine the relationship between oxygenation rate and each of the four stabilities above. Again, *P* values were rectified by *p*.*adjust* function.

### The role of elements within Jacobian *J*

Because warming/deoxygenation impact the dominant Lyapunov exponent LE of each four variables (ecological process, evolutionary process, whole community dynamic, or eco-evolutionary feedback), by changing the elements within the sequential Jacobian product 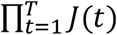 in eqn 4, we computed geometric mean of all elements within the Jacobian product as an indicator (hereafter called “mean strength”). Specifically, we computed the *mean strength of whole community*, by using the whole the Jacobian product in eqn 4, because the whole the Jacobian product stands for whole community dynamics. Similarly, we computed *mean strength of ecology*, by only using block matrix *A* to calculate its Jacobian product, because block matrix *A* stand for ecological process. The *mean strength of evolution* was computed, by only using block matrix *D* to calculate its Jacobian product, because block matrix *D* stand for evolutionary process. For *mean strength of eco-evolutionary feedback*, we aimed to compute how the eco-evolutionary feedback change mean *strength* of whole community, i.e. the difference between the *mean IS having the feedback* and the *mean strength not having the feedback*. Thus, we first set *B* and *C* in eqn 5 as zero (i.e. setting antidiagonal as zero), to get the *mean strength not having the feedback*. When not setting antidiagonal as zero, we can easily get the *mean strength having the feedback*. Finally, *mean strength of eco-evolutionary feedback*, was computed as the difference between the two, *i*.*e. mean strength having the feedback* subtracting *mean strength not having the feedback*.

Next, we employed linear regression to examine the effects of warming/deoxygenation on each of the four variables (*mean strength of whole community, ecology, evolution, or eco-evolutionary feedback*). Then we again employed linear regression to examine the association between *mean strength of whole community* and LE of whole community, the association between *mean IS of ecology* and LE of ecology, the association between *mean strength of evolution* and LE of evolution, and the association between *mean strength of eco-evolutionary feedback* and LE of eco-evolutionary feedback. Again all P values were rectified by *p*.*adjust* function.

## RESULTS

In simulated herbivore-resource communities, eco-evolutionary feedback destabilized community dynamics, as they increased the Lyapunov exponent (LE) (Fig 1d). Across 11 natural aquatic communities, these communities were on the edge of chaos (LE=0) (Fig 2). Ecological dynamics (i.e. abundance changes) stabilized over time, as LE were negative (Fig 2). Similarly, trait evolution stabilized over time (Fig 2). In contrast, eco-evolutionary feedback destabilized whole community dynamics, as it increases LE (Fig 2).

**Figure 1.**
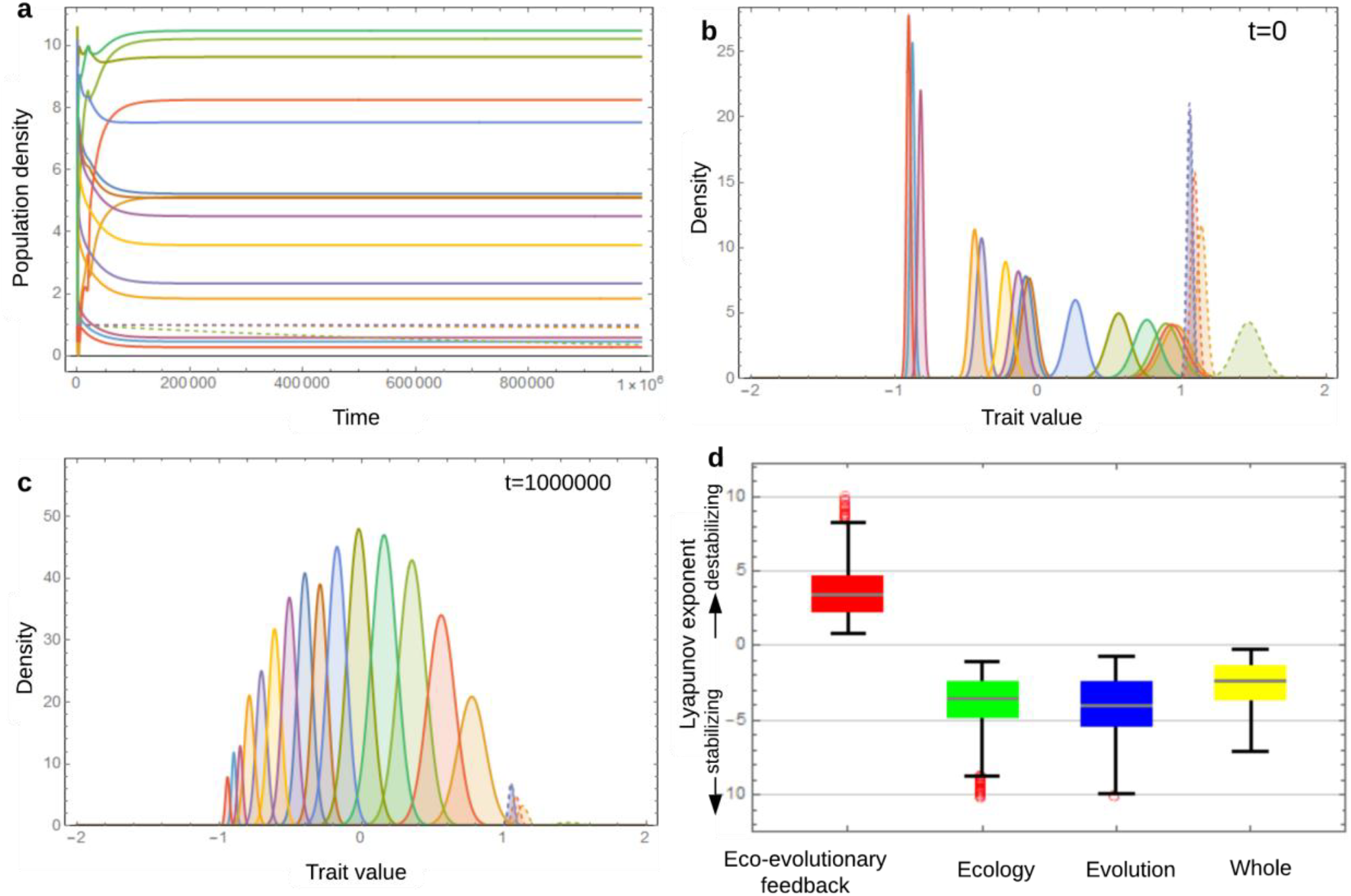
Population abundance, trait, and dominant Lyapunov exponent LE in simulated communities. **a.** population abundance over simulation (t=0 to 1000000). **b**. trait distribution at beginning (t=0). **c**. trait distribution at end of simulation (t=1000000). In **a-c**, solid lines indicate the producers, while the dashed lines indicate herbivores. **d**. dominant Lyapunov exponent LE for the four variable (eco-evolutionary feedback, ecological process, evolutionary process, whole community dynamic). In **d**, positive values for LE of eco-evolutionary feedback indicate that it destabilizes whole community dynamics over time, and vice versa. If LE of each the three variable (whole community dynamic, ecological process, evolutionary process) is lower than zero, indicating that it is stabilized over time, and vice versa.

**Figure 2.**
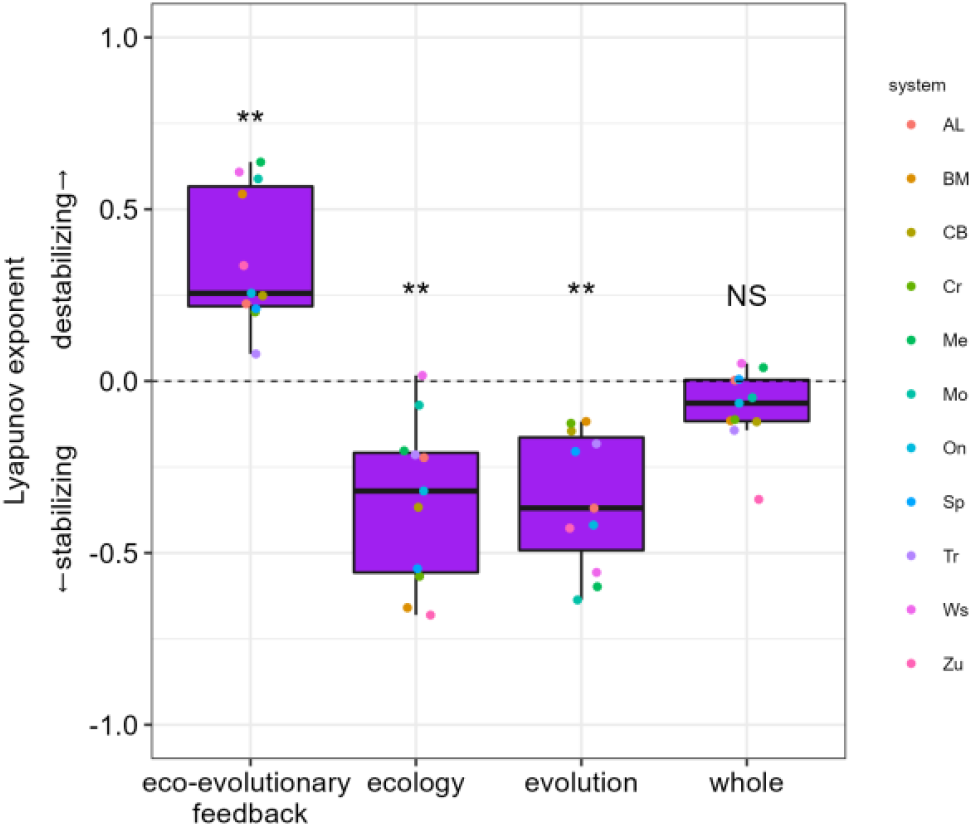
The dominant Lyapunov exponent LE of the eco-evolutionary feedback, ecological process, evolutionary process, and whole community dynamic. If LE of eco-evolutionary feedback is significantly higher than zero, indicating that it destabilized whole community dynamics over time, and vice versa. If LE of each the three variable (whole community dynamic, ecological process, evolutionary process) is significantly lower than zero, indicating that it is stabilized over time, and vice versa. The significance is examined by One-sample wilconxon test, and then be rectified by *p*.*adjust* function to adapt the facts that significance threshold (*P*=0.05) might be lenient. (\**P*<0.05, ** *P*<0.01, *** *P*<0.001)

Warming temperature destabilized the ecological process and made eco-evolutionary feedback even more destabilizing, while warming stabilized trait evolution (Fig 3a). We didn’t find warming had an effect on whole community dynamics.

**Figure 3.**
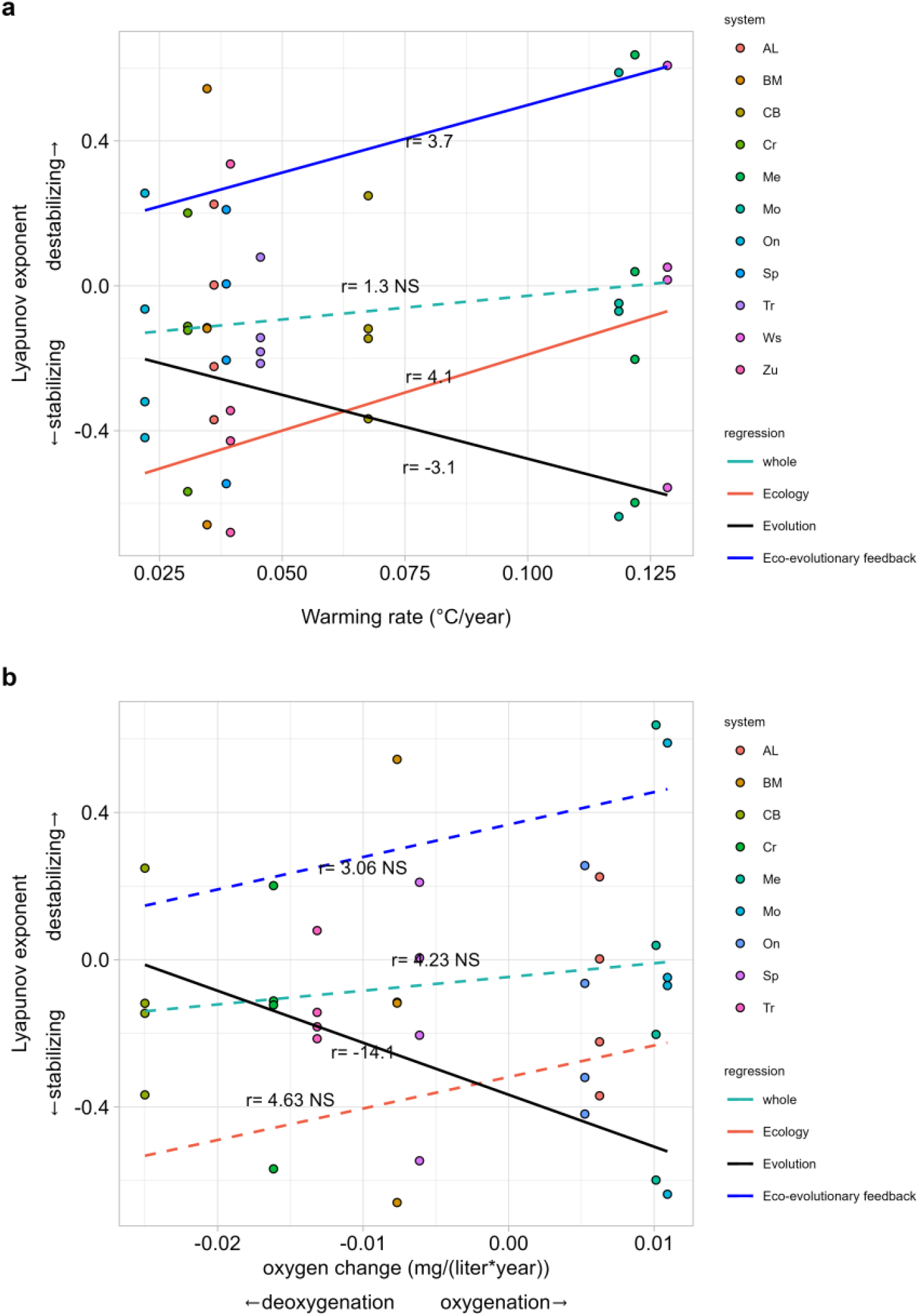
The effect of warming rate (a) and deoxygenation (b) on dominant Lyapunov exponent LE of eco-evolutionary feedback, ecological process, evolutionary process, and whole community dynamic. Linear regression was employed to test the association between warming rate and LE of each four variables (eco-evolutionary feedback, ecological process, evolutionary process, and whole community dynamic), and association between deoxygenation and LE of each four variables. A smaller LE indicates higher stability. The significance is rectified by *p*.*adjust* function.

Deoxygenation stabilized trait evolution (Fig 3b), while it had no effect on whole ecological process, eco-evolutionary feedback, and community dynamics (Fig 3b). The effects of warming and deoxygenation on stability above might be understood by change of the elements within Jacobian, which was used to compute LE.

Warming destabilized the ecological process and induced eco-evolutionary feedback more destabilizing, by increasing the mean strength (i.e. geometric mean of all elements within the Jacobian product) (Fig S1a). Higher mean strength was associated with higher LE (Fig S1c). In contrast, warming stabilized trait evolution of these communities by decreasing mean strength (Fig S1a). In addition, deoxygenation destabilized trait evolution, as deoxygenation was marginally associated with higher mean strength (Fig S1b). Finally, we found that length of time series or size of community matrix had no effects on LE (Fig S2).

## DISCUSSION

We found that the eco-evolutionary feedback destabilizes the community dynamics in both simulated and natural communities. In natural aquatic systems, warming made the eco-evolutionary feedback more destabilizing. In addition, trait evolution stabilized itself over time while warming and deoxygenation made the evolution stabilizing and destabilizing, respectively.

Previous modelling evidences showed eco-evolutionary feedback can stabilize predator-prey communities to stable state (Martis 2022) or can trigger a three species food chain transition from a stable state to be unstable (Tanaka 2022). In addition, experimental evidence also showed a host–virus system (Frickel *et al*. 2016) and cooperative species (Sanchez & Gore 2013) with demonstrated eco-evolutionary feedback converge to steady state, whereas a herbivore-resource (i.e. a rotifer-algae) system didn’t converge to steady state but exhibit cycles (Yoshida *et al*. 2003, 2007; Haafke *et al*. 2016). However, these evidence is from either simple modelling simulations and/or a small number of species. We lack evidence for species rich systems. Here we showed that eco-evolutionary feedback destabilized community stability in both simulated and natural species-rich systems.

In addition, we found warming led to the eco-evolutionary feedback more destabilizing. Empirical evidence (Cherabier & Ferrière 2022) and schematic reviews (Faillace *et al*. 2021) highlighted that warming could change the magnitude of eco-evolutionary feedback. For example, Cherabier and Ferrière (2022) showed warming can decrease primary production via mediating the eco-evolutionary feedback in nutrient-poor environments. However, no evidence documents how warming impact the effect of the feedback on community stability. We found that warming resulted in the effect of the feedback on stability more unstable, by increasing the mean strength (i.e. geometric mean of all elements within the Jacobian product).

Deoxygenation could change ecological process and trait evolution. For example, lower oxygen level promoted the growth of cyanobacteria, which led to phytoplanctivorous invertebrate more tolerant to dietary cyanobacteria. However, no study examined how deoxygenation change trait evolution in species-rich natural systems. We showed deoxygenation led to the trait evolution destabilizing, which might be caused by increasing the mean strength. We didn’t know exact mechanism of how deoxygenation increasing the mean strength. One potential mechanism driving this result is that natural selection may favor deoxygenation-tolerant species. As its offspring becomes deoxygenation-tolerant, consumption rate and net growth rates probably increase as they adapt to deoxygenation.

We admit the limitation in our study. We only consider a single trait for each species. However, species in nature can have multiple heritable traits. In addition, to simplify the modelling simulation, we assume the focal species under selection has only a single allele at the focal locus. However, a selection can be govened by two or more alleles.

Overall, we have shown the eco-evolutionary feedback could affect community stability, and how climate changes the feedback in natural aquatic systems. Our results suggest that a combination of eco-evolutionary feedback and climate change may help us to better understand the ecological and evolutionary changes in nature.

## AUTHORSHIP

QHZ, FDL and GB designed the research. QHZ developed the models and analyzed the data; QHZ, FDL and GB drafted the manuscript.

## ACKNOWLEDGEMENTS

We thank all participants in each long-term monitoring site. Q.Z. acknowledges funding from the concerted research action (ARC) from the special research fund from the University of Namur, Belgium (ARC grant DIVERCE, Convention 18/23-095).

## DATA ACCESSIBILITY STATEMENT

The data and codes supporting the results are archived at GitHub (https://github.com/QZhao16).

## SUPPLEMENTARY FIGURES

**Figure S1.**
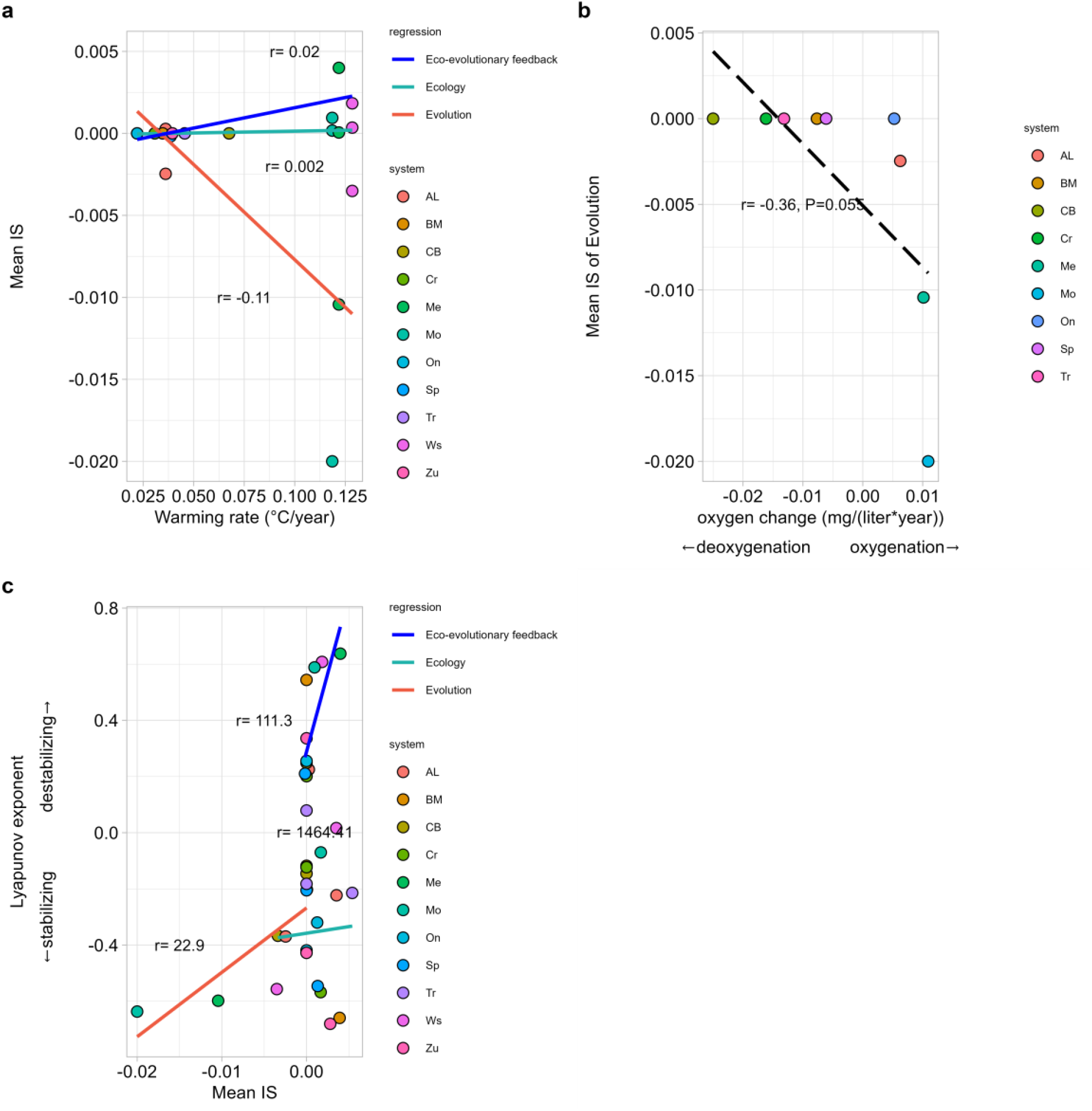
The effect of warming rate on mean interaction strength (mean IS) of eco-evolutionary feedback, ecological process, evolutionary process (a), the effect of deoxygenation on mean IS of evolutionary process (b), and the effect of mean IS of each variable (eco-evolutionary feedback, ecological process, or evolutionary process) on its dominant Lyapunov exponent LE (c). Linear regression was employed to test the association between them. The significance is rectified by *p*.*adjust* function. The solid lines indicate significant effects, and no regression lines indicate nonsignificant effects. A marginal significance is marked (**b**).

**Figure S2.**
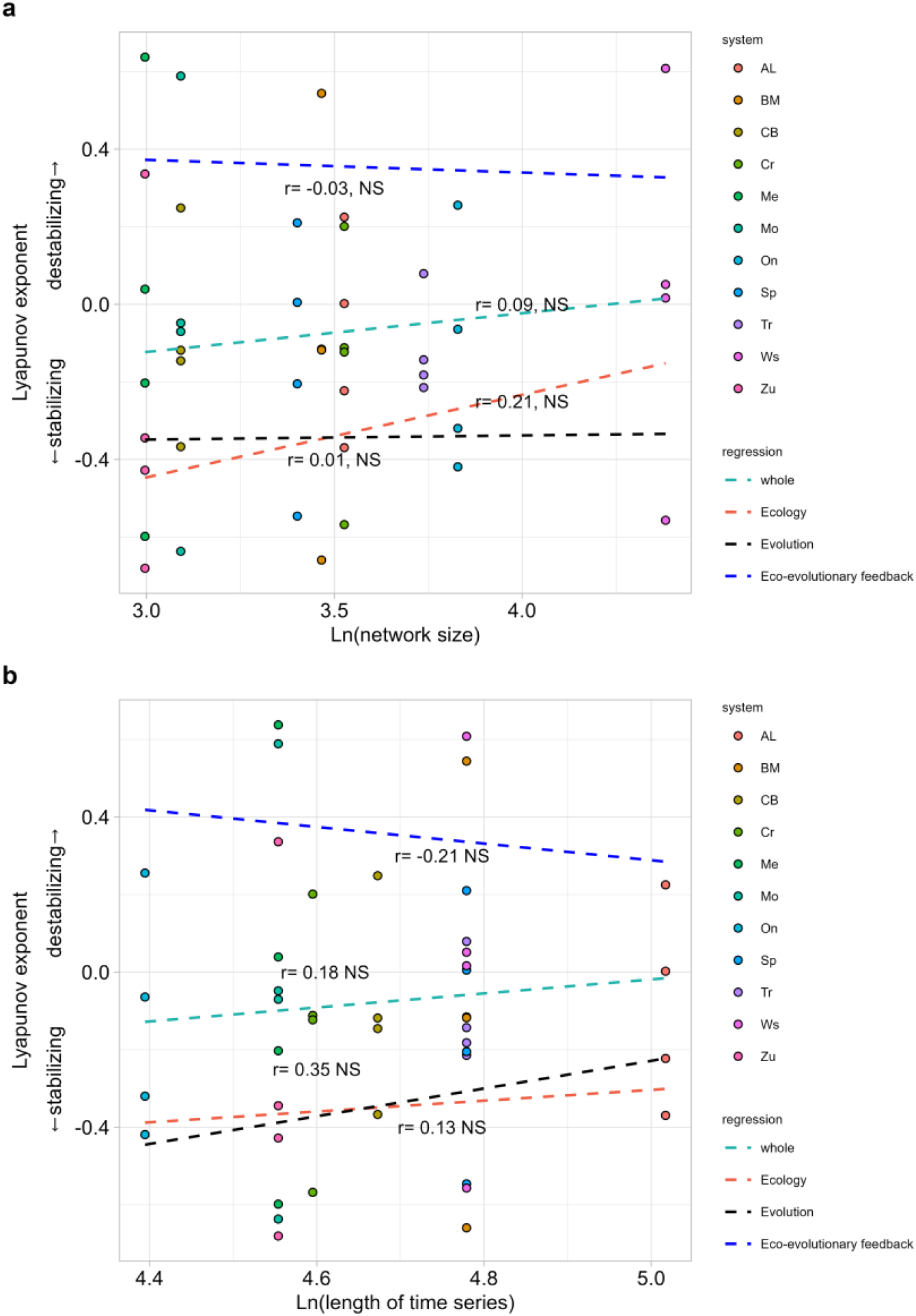
The effect of network size (a) and length of time series (b) on dominant Lyapunov exponent LE of eco-evolutionary feedback, ecological process, evolutionary process, or whole community dynamic dominant. Linear regression was employed to test the association between network size and LE of each four variables (eco-evolutionary feedback, ecological process, evolutionary process, or whole community dynamic), and association between length of time series and LE of each four variables. A smaller LE indicates higher stability. The significance is rectified by *p*.*adjust* function. The solid lines indicate significant effects, and no regression lines indicate nonsignificant effects.

**Figure S3.**
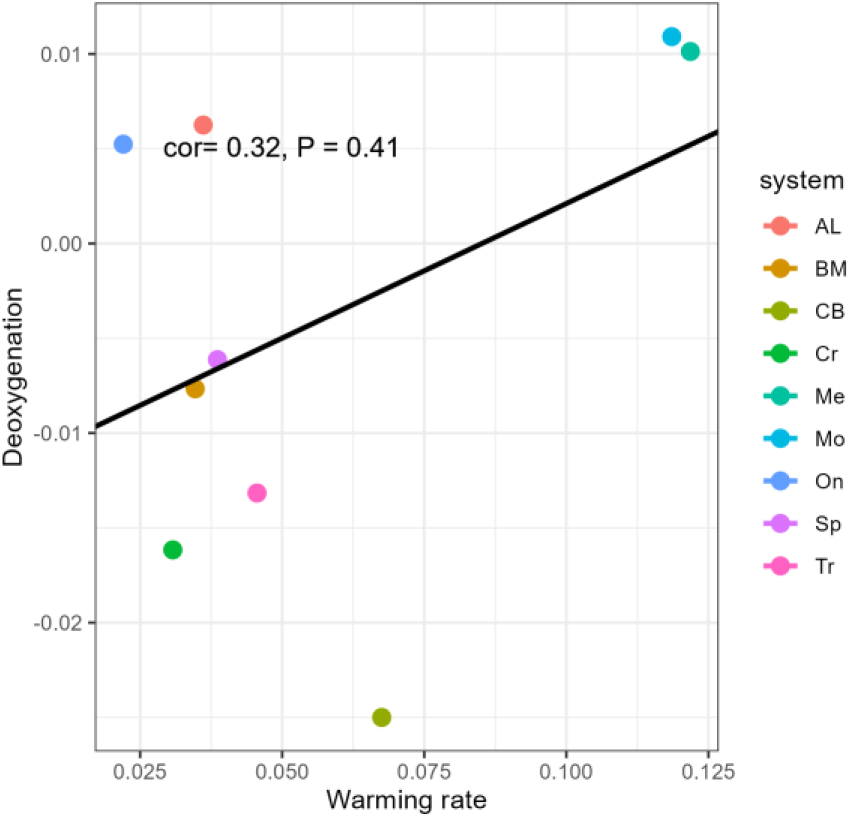
Correlation between warming and deoxygenation. The correlation was examined by spearman correlation. Nine system was used to analysis, because Lake Zurich (Zu) and Wadden sea (Ws) didn’t report dissolved oxygen concentration.

